# A coarse-graining, ultrametric approach to resolve the phylogeny of prokaryotic strains with frequent homologous recombination

**DOI:** 10.1101/094599

**Authors:** Tin Yau Pang

## Abstract

A frequent event in the evolution of prokaryotic genomes is homologous recombination, where a foreign DNA stretch replaces a genomic region similar in sequence. Recombination can affect the relative position of two genomes in a phylogenetic reconstruction in two different ways: (i) one genome can recombine with a DNA stretch that is similar to the other genome, thereby reducing their pairwise sequence divergence; (ii) one genome can recombine with a DNA stretch from an outgroup genome, increasing the pairwise divergence. While several recombination-aware phylogenetic algorithms exist, many of these cannot account for both types of recombination; some algorithms can, but do so inefficiently. Moreover, many existing algorithms require that a substantial portion of each genome has not been affected by recombination, a sometimes unrealistic assumption. Here, we propose a novel coarse-graining approach for phylogenetic reconstruction (CGP), which is recombination-aware, applicable even if all genomic regions have experienced substantial amounts of recombination, and can be used on both nucleotide and amino acid sequences. CGP considers the local density of substitutions along pairwise genome alignments, fitting a model to the empirical distribution of substitution density to infer the pairwise coalescent time. Given all pairwise coalescent times, CGP reconstructs an ultrametric tree representing vertical inheritance. Based on simulations, we show that the proposed approach can reconstruct ultrametric trees with accurate topology, branch lengths, and root positioning. Applied to a set of *E. coli* strains, the reconstructed trees are most consistent with gene distributions when inferred from amino acid sequences, a data type that cannot be utilized by many alternative approaches.

**AUTHOR SUMMARY:** In homologous recombination, segments of foreign DNA overwrite similar segments of a prokaryotic genome. A single recombination event can simultaneously introduce many DNA substitutions. This disturbs phylogenetic signals, making it difficult to reconstruct prokaryotic family trees. While a handful of recombination-aware phylogenetic algorithms have been proposed, most do not take all effects of recombination into account; others rely on the frequently unrealistic assumption that a substantial part of a genome has not been affected by recombination at all. Here, we introduce a novel approach to phylogenetic reconstruction, which estimates the age of the most recent common ancestor of two strains from the density distribution of DNA or amino acid substitutions between their genomes. The proposed phylogenetic tree is the tree most compatible with these age estimates. Based on nucleotide or amino acid sequences, our approach accurately predicts the topology, branch lengths, and root positioning of prokaryotic family trees.

## INTRODUCTION

The transfer of DNA stretches from one prokaryotic genome to anotherotermed horizontal gene transfer (HGT) or lateral gene transfer (LGT)–is a major driver of prokaryotic evolution [1]. It is caused by a variety of mechanisms, including transformation, transduction, conjugation, and gene transfer agents [2,3]. Many prokaryotic genomes encode defense systems against foreign DNA, such as the restriction modification system [4]. A foreign DNA stretch that enters the prokaryotic cell and survives these host defenses may be incorporated into the host genome. If the incoming DNA stretch is highly similar to a stretch on the host genome, homologous recombination may occur, where the incoming DNA stretch homologously recombines with the host stretch and overwrites it [5,6]. Alternatively, the incoming stretch may be inserted directly into the host genome through non-homologous recombination.

HGT allows the fast spread of genes through prokaryotic pangenomes, and facilitates rapid adaptation to environmental changes. A point in case is the spread of antibiotic resistance genes in pathogenic bacteria via HGT [7]. But recombination is also crucial for the long-term maintenance of prokaryotic populations, as it helps to repair DNA damaged by deleterious mutations, thereby avoiding the mutational meltdown of Muller’s ratchet [8]; in that sense, prokaryotic recombination may fulfill the same function as does sex in eukaryotes. Computational modelling also suggests that recombination may help prokaryotes to purge selfish mobile genetic elements [9].

Recombination can severely affect phylogeny reconstructions. Its effects on genome divergence are complex. Depending on the circumstances, recombination can speed up the divergence of a genome pair or slow it down [6]; its effects may severely affect the accuracy of estimated branch lengths of phylogenetic tree. For example, (i) when a stretch of genome X is replaced by DNA from genome Y, some of the single nucleotide polymorphisms that previously differentiated X and Y will be erased, shortening the apparent evolutionary distance between the two genomes. Conversely, (ii) when X recombines with a DNA stretch of an outgroup genome (a genome that diverged before the split of the X and Y lineages), then this introduces further nucleotide polymorphisms into X, thereby increasing the apparent X-Y distance.

Multilocus sequence typing (MLST) aims to extract sequences of housekeeping genes from prokaryotic genomes, which can then be utilized to resolve evolutionary relationships [10]. However, MLST genes may also experience frequent recombination, and phylogenetic reconstruction without accounting for recombination can compromise the resulting trees [11]. In fact, the frequency with which recombination affects a gene can be of the same order of magnitude as the corresponding mutation rate [5]. If two lineages recombine with each other, application of conventional phylogenetic algorithms without accounting for recombination will generally lead to an underestimation of the age of the common ancestors [12]. When there are more than two strains, recombination affects the reliability of inference of relative divergence times between strains and may hence compromise both tree topology and branch length estimates. There are several popular recombination-aware algorithms, including ClonalFrameML (and its predecessor ClonalFrame) [13,14], the Bacter package in BEAST2 (which implements the ClonalOrigin model) [15,16], and Gubbins [17]; there are also non-phylogenetic algorithms that detect recombination, such as BratNextGen and fastGEAR [18,19]. While these algorithms can identify genomic stretches with high numbers of substitutions due to recombination with distant strains and thus account for type (ii) recombination effects, many do not take type (i) recombination effects into account; Bacter can account for type (i) effects, but is not computationally efficient for long genome sequences. Some of these algorithms rely on the assumption of low recombination rates, such that a substantial part of a genome remains clonal and has not been affected by recombination. This is unrealistic at least for some bacteria: *e.g.*, *E. coli* strains whose DNA sequences have diverged by more than 1.3% share very few segments larger than a few kilo-basepairs that have not been affected by recombination in at least one of the two lineages [5].

In this paper, we present a novel model that does not assume low recombination rates. This model follows a coarse-graining approach, which considers the local density of substitutions on a sequence alignment instead of site-specific substitutions, and which is applicable to both nucleotide and amino acid sequences. The source code implementing this model is available at https://github.com/TinPang/coarse-graining-phylogenetics.

## MATERIALS AND METHODS

### The CGP model describing the evolution of single site polymorphism distributions of genome pairs in a Fisher-Wright population

Our coarse-graining phylogenetic (CGP) model is based on a Fisher-Wright haploid population with nonoverlapping generations, constant population size, and homologous recombination [20,21]. In this framework, a node in one generation inherits the genome of a random node in the previous generation, followed by mutation and homologous recombination. The CGP model considers segments instead of nucleotide / amino-acid sites as the basic unit of a genome, because the local density of single site polymorphism (SSP) can be defined conveniently on segments; an SSP can be a single nucleotide polymorphism (SNP) on a nucleotide sequence, or a single amino acid polymorphism (SAP) on an amino acid sequence. A genome sequence is divided into *L*_*seg*_ consecutive and non-overlapping segments, where every segment has length *ls* (*i.e.*, consists of *l*_*s*_ sites). The rate of mutation is *μ* per site per generation. The rate for a site to be covered by a foreign DNA stretch attempting to recombine with the host genome is *ρ* per generation; the rate for a segment to be covered by a recombination-attempting DNA stretch is also approximatly *ρ*, assuming that the segment is much shorter than the DNA stretch. Here, *ρ*=*ρ*_*ini*_*L*, where *ρ*_*ini*_ is the probability for a recombination-attempting foreign DNA stretch to start at any given site, and *L* is the average length of the foreign DNA stretch. When a recombination attempt happens on a segment, it either succeeds, and the foreign DNA replaces the host DNA at the segment, or it fails. The success rate of an attempt is approximately exp(-*δ*/*δ*_TE_),where *δ* is the divergence between the incoming DNA and the host DNA, and *δ*_TE_ is the transfer efficiency, a constant that governs the success rate [5]. The average site divergence in the population is denoted as *θ*, with *θ*=2*μN*_*e*_ and population size *N*_*e*_.

CGP’s model [5,6] considers the evolution of a SSP distribution between a pair of genomes, X and Y. As the alignment of genome X and Y is divided into *L*_*seg*_ consecutive and non-overlapping segments with *ls* sites, let *f*(*x*|*t*) be the distribution of segment divergence, where *x*=0, 1, _*ls*_ represents the number of SSPs on a segment of the XY alignment, *t*≥0 is the (continuous) XY coalescent time, and *f*(*x*|*t*) is normalized to unity by summing over *x*. To save computational resources, we assume an upper bound 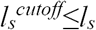 to *x*. At *t*=0, the most recent common ancestor (MRCA) of XY splits into two lineages; initially, the two have identical genomes, and thus *f*(*x*|0)=δ*x*,0 (where δ*x*,0 is the Kronecker delta, *i.e.*, *f*(*x*|0) is non-zero only at *x*=0). At *t*>0, mutations and recombinations occur, and the evolution of *f*(*x*|*t*) is described by the following equation:

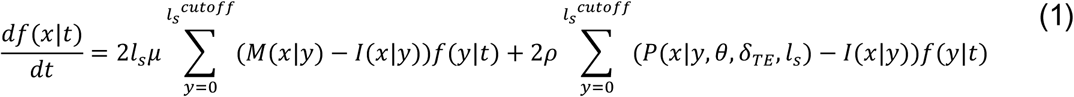

The first term of Eq (1) accounts for mutations on a segment—M(*x*|*y*) models a mutation event, where a segment in the pair XY with *y* SSPs increases to *x*=*y*+1 SSPs during a mutation (*i.e.,* M(*x*|*y*)=0 for *x*≠*y*+1); I(*x*|*y*) is the identity matrix. For simplicity, we ignore back mutations.

The second term accounts for recombination—P(*x*|*y*,*θ,δ*_*TE*_,*l*_*s*_) models a recombination event (see Eq. (S4) of *Dixit et al.* [5] or Eq. (3) of *Dixit et al.* [6] for a detailed derivation). Since a segment can recombine with its counterpart on another genome, it assumes that each segment of a genome, along with its counterparts in different genomes of the population, have their own phylogeny, which is detached from the genomes phylogeny, and the segment population structure is approximated by the coalescent model. For an attempted recombination between Y and an external donor D, we can use the coalescent model to calculate the probability distribution for the segment divergence *δ* between D and X, and then obtain *x* from *x*=*l*_*s*_*δ*. As mutation and recombination can equally occur on either X or Y, there is a factor 2 attached to both terms. See Supplementary Text for the exact form of P(*x*|*y*,*θ,δ*_*TE*_,*l*_*s*_). We solved Eq. (1) with boundary condition *f*(*x*|0)=δ*x*,0 to obtain the theoretical SSP distribution *f*(*x*|*t*) at different coalescent times *t*.

We fit the the theoretical distribution obtained with the CGP model to an empirical SSP distribution to infer the coalescent time of the pair of genomes. Let us consider an alignment for a genome pair XY that is divided into *L*_*seg*_ segments, with empirical SSP distribution *g*XY(*x*) following the normalization condition:

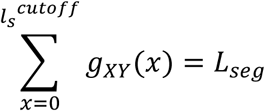

Let us denote the theoretical distribution as *f*_*μ,ρ,θ,δte*_*x*|*t*), which is normalized to unity The probability to observe the empirical distribution *g*XY(*x*) given the theoretical distribution *f*_*μ*_,_*ρ,θ,δTE*_(*x*|*t*) is

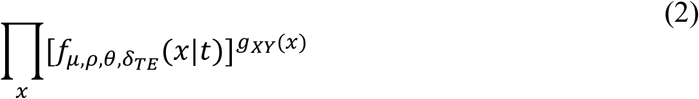

If we take the logarithm of this expression, it becomes the (negative) cross entropy between *g*(*x*) and *f*_*μ,ρ,θ,δTE*_(*x*|*t*) [22,23]. The higher their similarity, the higher is this negative cross entropy; it attains its maximum when *f*_*μ,ρ,θ,δTE*_(*x*|*t*) is equal to *g*(*x*).

Suppose that we have *n* genomes (X_*i*_, *i*=1,…,*n*), where their phylogeny is described by an ultrametric tree *T*; the *n*(*n*-1)/2 pairwise SSP distributions have evolved according to the model with parameters *μ, ρ, θ,* ^*δ*^*TE*. Let *tT*(X_*a*_,X_*b*_) be the coalescent time of X_*a*_ and X_*b*_ in the tree *T*. We use a score function, *S*(X_1_,X_2_, o,X*n*|*μ*, *ρ*, *θ*, ^*δ*^TE,*T*), which is defined as the logarithm of the probability to observe the *n*(*n*-1)/2 empirical SSP distributions given the model and the tree, to quantify the model fit to the empirical SSP distributions. This score is the summation of the *n*(*n*-1)/2 negative mutual entropy terms:

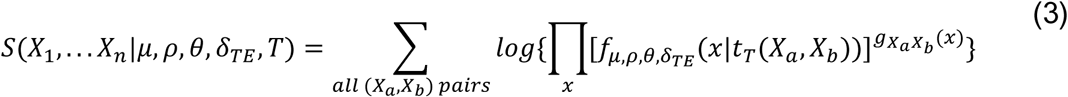

Since the *n*(*n*-1)/2 SSP distributions are not completely independent of each other, Eq. (3) is not exactly a probability and so we call it a score. We developed an algorithm that samples the tree+model space and searches for the configuration with the maximum score using Monte Carlo simulation with annealing and Metropolis acceptance (See Supplementary Text for details).

## RESULTS

### A coarse-graining approach to phylogenetic reconstruction

The coarse-graining phylogenetics (CGP) model is based on a mathematical model [5,6] that quantitatively describes the evolution of genomic sequence divergence; this model is applicable to both nucleotide and amino acid sequences, and does not assume low recombination rate. Recombination can introduce DNA stretches characterised by a high density of substitutions, and the CGP model considers substitution densities defined on the genomic segments. A nucleotide genome sequence alignment (or a corresponding concatenation of amino acid sequence alignments) is divided into a chain of consecutive, non-overlapping segments, each with *l*_*s*_ sites; for a pair of genomes, the single site polymorphisms (SSPs) on each segment are counted, resulting in an SSP distribution. CGP takes the SSP distribution of every pair of considered genomes as input. The coalescent time of two genomes can be inferred by fitting the CGP model to the empirical SSP distribution. The ultrametric tree describing the vertical component of inheritance among *n* genomes can be inferred from the coalescent times resulting from the fits to the *n*(*n*-1)/2 empirical SSP distributions, implemented by the score function of Eq. (3). We developed the CGP algorithm, which employs Monte Carlo simulation to sample the model+tree space, identifying the tree and parameters that result in the highest score.

### Testing the CGP algorithm on simulated genomes

We performed Fisher-Wright simulations with recombination to generate genome sequences, allowing us to test different phylogenetic reconstruction algorithms. In the simulation, each recombination-attempting DNA stretch starts at a random site of a genome, with equal chance to be either a micro (geometric distribution, mean 100bp) or a macro stretch (geometric distribution, mean 1kb). We used three sets of parameters that correspond to prokaryotic populations with r/m=2, 40, 80 (r/m is the ratio between substitutions contributed by mutations and by recombinations; these three settings are denoted as low, intermediate, and high recombination level, respectively), and prepared the test groups, each with 4-10 genome sequences. For comparison, r/m values observed in nature range from 0.02 to 63.6 [11].

The MRCA of a group of random genomes in a simulated population has an average age close to the age of the population root node *t*_root_. We would like to mimic the condition where a single lineage diverges from the rest of the population and forms its own subpopulation, so that the genomes in its subpopulation continue exchanging DNA among themselves and with the rest outside. Hence, when picking genomes in the population to form test groups, we constrained the age of the MRCA of the genomes in a test group, *t*_test-group-root_, to be *t*_test-group-root_≪_*t*root_ (see Supplementary Text for the details and Supplementary File S3 for the genome sequences in each test-group).

We applied CGP, as well as the previously published methods RaxML [24], ClonalFrameML [13], and Gubbins [17] to the sequences of each test group. The RAxML and Gubbins trees are midpoint-rooted. ClonalFrameML requires an initial tree as input and we used the RAxML tree. CGP uses segment size *l*_*s*_=150 and *l*_*s*_^*cutoff*^=100. We compared each reconstructed tree with the true tree, measuring their unrooted symmetric distance (SD) [25], as well as their rooted and unrooted branch score distance (BSD) [26] to quantify the accuracy of the reconstructed phylogeny (see Supplementary File S1 for the these values); the lower the unrooted SD / unrooted BSD / rooted BSD, the more accurate is the topology / branch lengths / root positioning, respectively. We normalized the branch lengths of each tree by its total branch length when calculating BSD.

CGP can predict the topology of a phylogeny of vertical inheritance as accurately as the other algorithms. Figure 1 shows the histograms of unrooted SD; ClonalFrameML is excluded as it uses the topology of RAxML trees. The distributions of SD of CGP are not significantly different from the distributions of RAxML and Gubbins. Two-sided Wilcoxon signed-rank tests (WSRT) at low, intermediate, and high recombination levels resulted in *p*=0.25, 0.69, 0.54 between CGP and RAxML, and *p*=0.25, 0.38, 0.92 between CGP and Gubbins.

**Figure 1.**
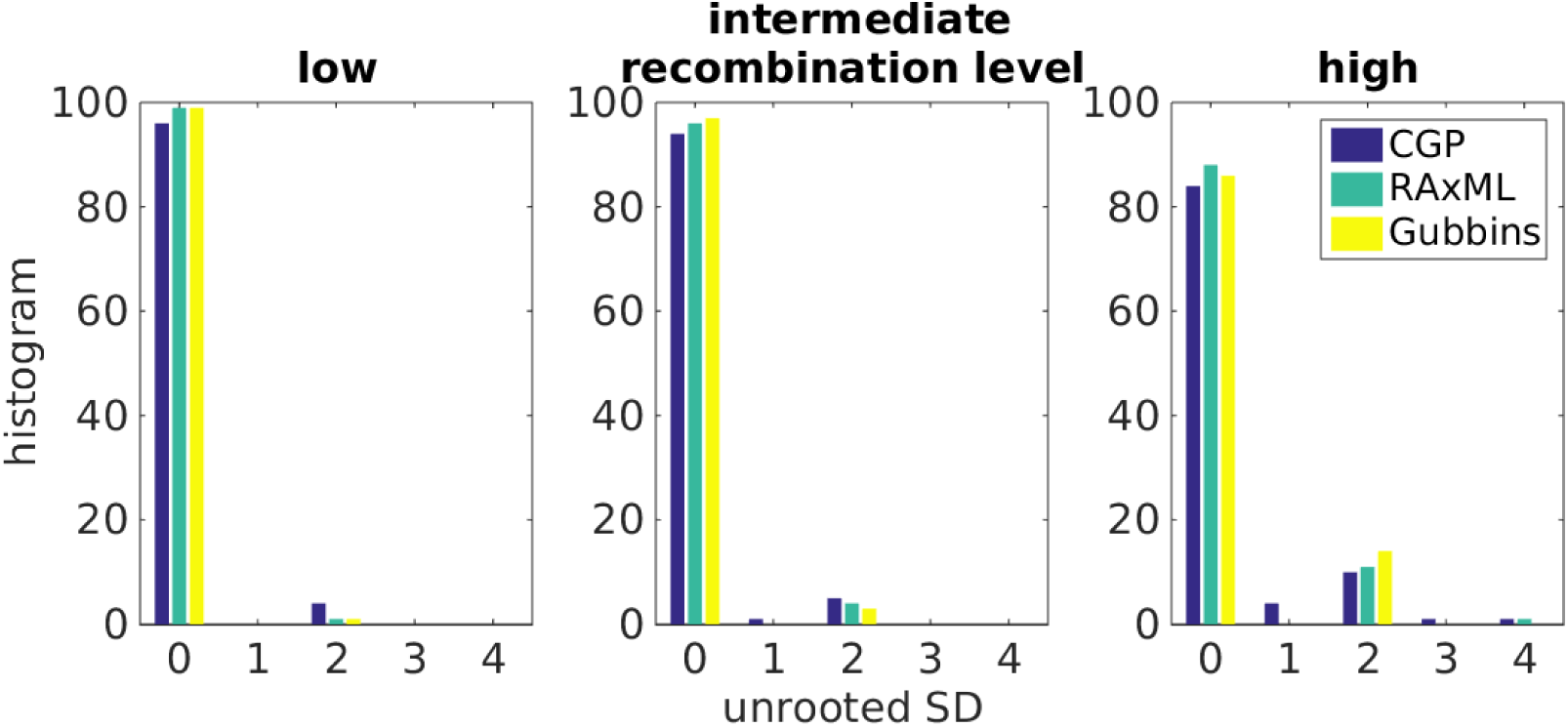
Histograms showing the distributions of the unrooted symmetric distance (SD) between true trees and tree reconstructed by CGP, RaxML, and Gubbins, from genome sequences derived from Fisher-Wright simulations with low, intermediate, and high recombination levels.

Branch length predictions are more accurate with CGP than with alternative programs at higher levels of recombination. Figure 2 plots the distributions of the unrooted BSD of different algorithms and their z-scores; the unrooted BSD of trees reconstructed by different algorithms on the same test group sequences are pooled together to calculate their z-scores to help data visualization. The distribution of the unrooted BSD of CGP is significantly lower than RAxML and Gubbins (*p*<10^-^10^^ at low, intermediate, and high recombination level compared to both RAxML and Gubbins). The unrooted BSD of CGP is significantly lower than ClonalFrameML except at low recombination levels (*p*=0.76, 2.2×10^-^7^^, 10^-^7^^ at low, intermediate, and high recombination levels).

**Figure 2.**
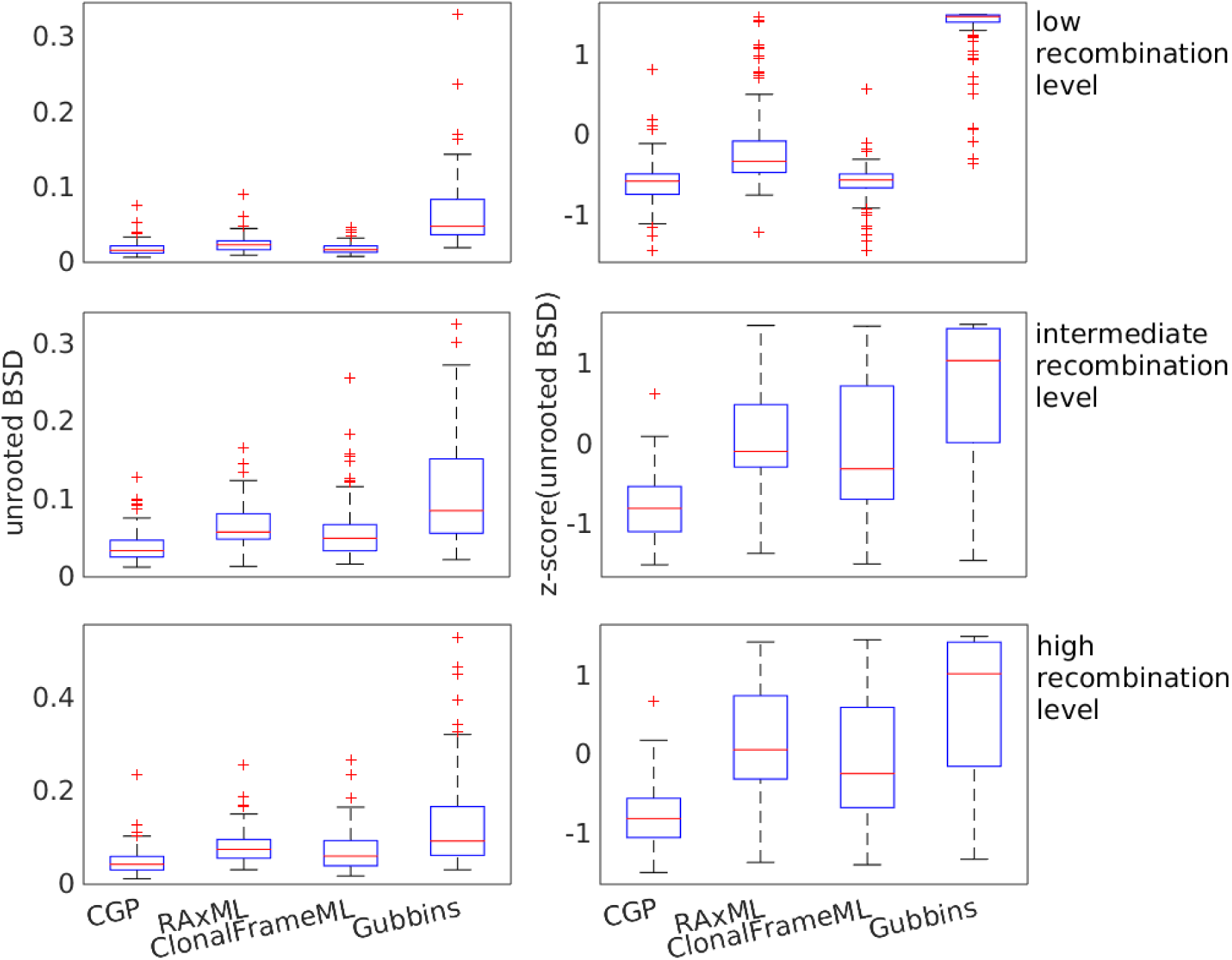
Boxplots showing the distributions of the unrooted branch score distance (BSD) and the distributions of the z-score of unrooted BSD between the true trees and trees reconstructed by CGP, RaxML, and Gubbins, from genome sequences from Fisher-Wright simulations with low, intermediate, and high recombination levels.

CGP can perform accurate root positioning. Figure 3 plots the distribution of the rooted BSD and their z-scores; the rooted BSD of trees reconstructed by different algorithms on the same test group sequences are pooled together to calculate their z-scores. The distribution of the rooted BSD of CGP is significantly lower than the other algorithms (*p*<2×10^-^17^^ at all recombination levels for CGP compared with the other three algorithms).

**Figure 3.**
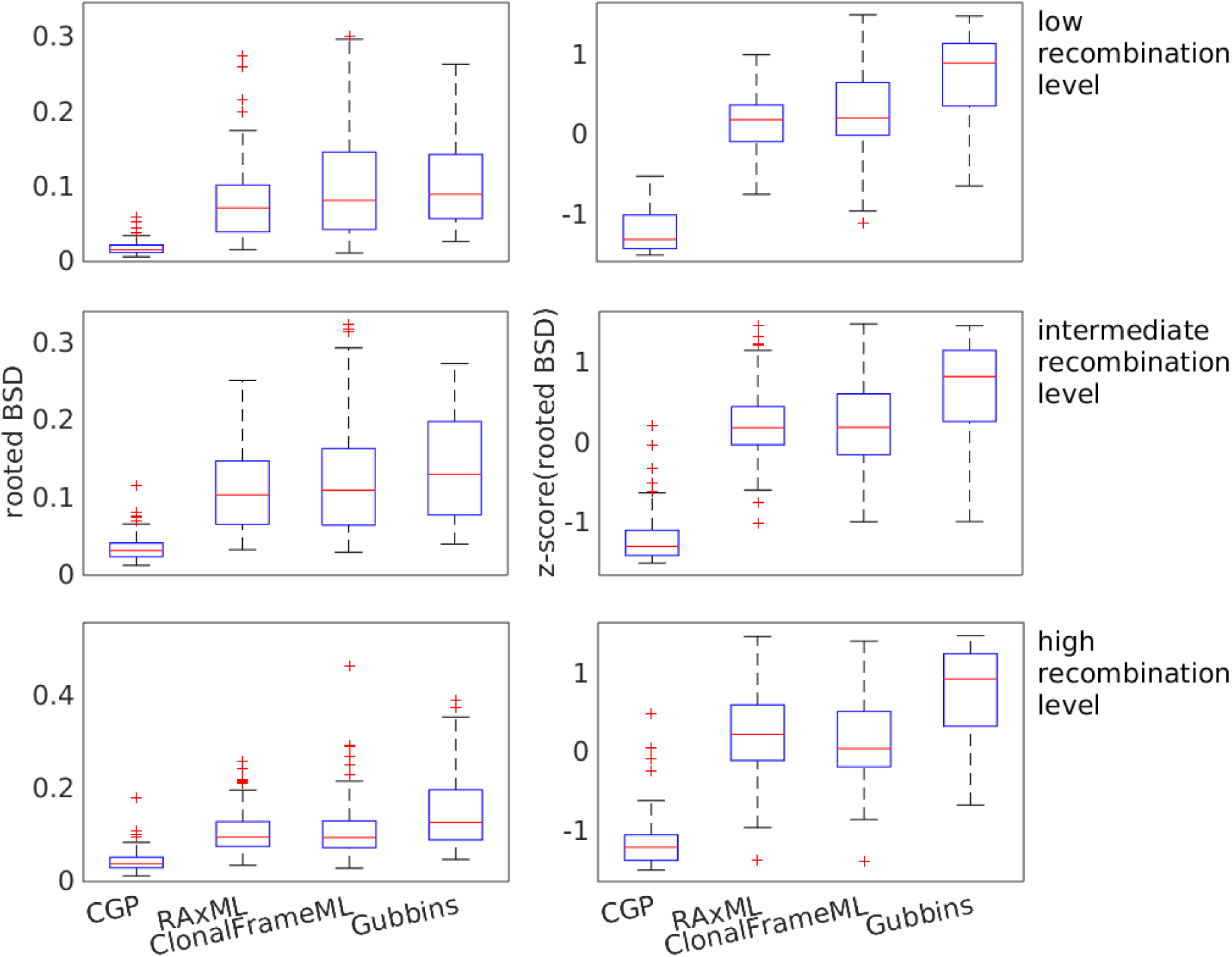
Boxplots showing the distributions of the rooted branch score distance (BSD) and the distributions of the z-score of rooted BSD between the true trees and trees reconstructed by CGP, RaxML, and Gubbins, from genome sequences from Fisher-Wright simulations with low, intermediate, and high recombination levels.

### Testing the CGP algorithm on real *E. coli* genomes

We tested CGP, RAxML, ClonalFrameML, and Gubbins using *E. coli* and *Shigella* genome sequences (see Supplementary Table S1 for their names); we refer to them as *E. coli*, as these two species have intertwined phylogenies. We prepared test groups, each with 10 random strains, where each strain is represented by a nucleotide and an amino acid sequence made from its concatenated core genes (see Supplementary File S2 for the strains in each test group, and also the 1,636 orthologous gene families of core genes; see Supplementary File S3 for their sequences). We applied CGP, RAxML, ClonalFrameML (with the topology from RAxML trees), and Gubbins on the nucleotide sequences, and CGP and RAxML on amino acid sequences; the RAxML and Gubbins trees are midpoint-rooted. CGP uses segment length *l*_*s*_=150 and *l*_*s*_^*cutoff*^=100 for nucleotide sequences, and *l*_*s*_=50, *l*_*s*_^*cutoff*^=50 for amino acid sequences.

To assess the accuracy of the phylogenetic trees reconstructed by the different algorithms, we compared the reconstructed trees with the phylogenetic signal inferred from the distribution of orthologous gene families in different genomes. We applied the GLOOME algorithm [27], which considers the interior nodes of the tree as ancestral strains and reconstructs their gene distribution; it takes a tree and the presence-and-absence of genes across the extant strains as input, and performs a reconstruction of presence-and-absence of genes in the ancestral strains based on the GLOOME posterior likelihood (GPL). We used GPL as a score to quantify the accuracy of the tree fed into GLOOME; the more consistent the phylogenetic signal from the gene distributions with a given tree, the higher the GPL (see Supplementary File S2 for the GPL values of the reconstructed trees).

Figure 4 plots the distribution of the GPLs and the corresponding z-scores; the GPLs of trees of the same test groups reconstructed by different methods are pooled together to calculate the z-scores. Trees reconstructed from amino acid sequences have a higher GPL than trees calculated from nucleotide sequences; moreover, CGP trees based on amino acids are more accurate than tress calculated using RAxML (*p*<4×10^-^14^^ when comparing CGP on amino acid sequences with any other algorithm). Considering only trees reconstructed from nucleotide sequences, the CGP trees generally have higher GPL than RAxML, ClonalFrameML, and Gubbins trees (*p*=5.2×10^-^4^^, 0.049, 1.4×10^-^15^^, respectively).

**Figure 4.**
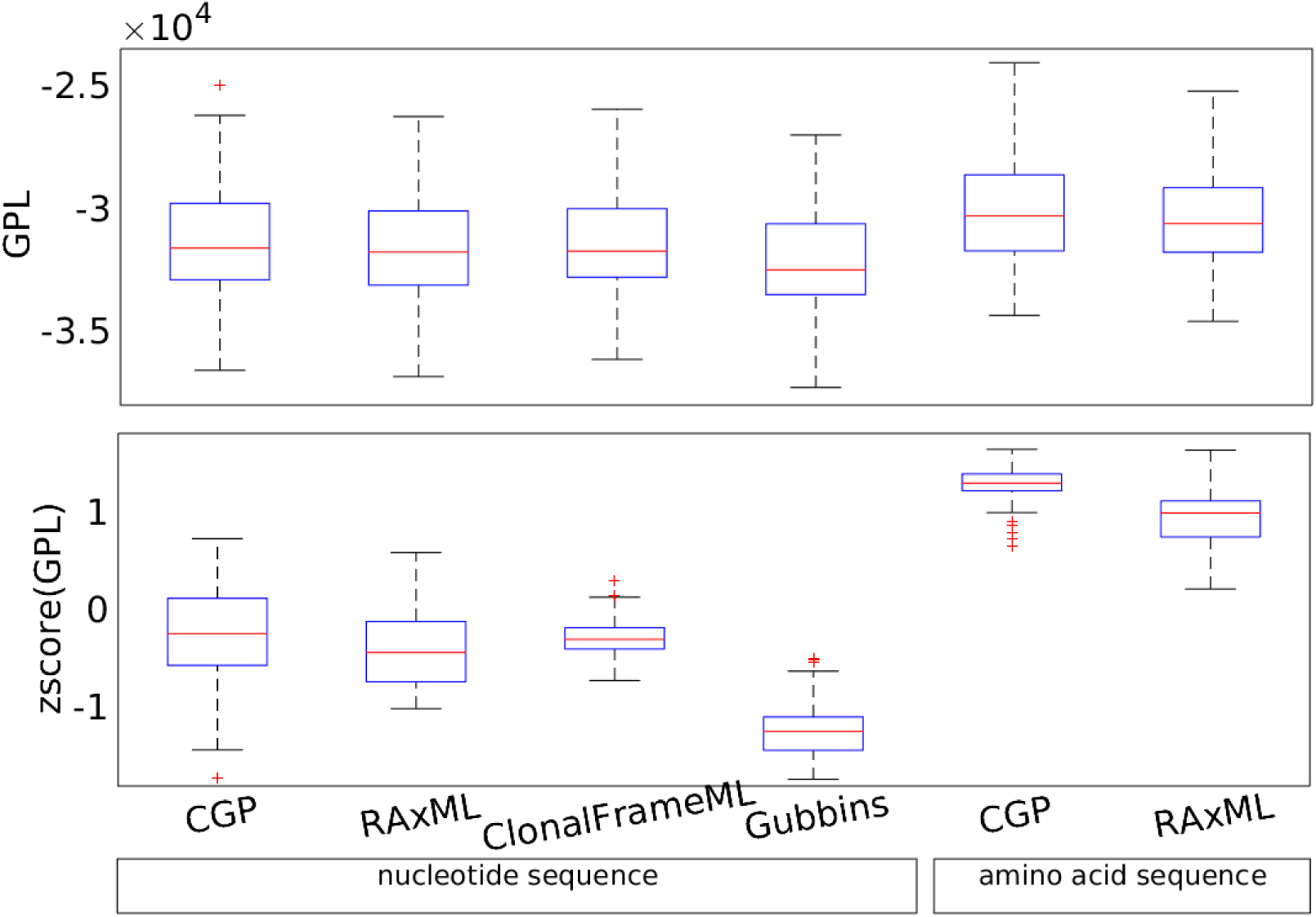
Boxplots showing the distributions of the GLOOME posterior likelihood (GPL) and the distributions of the z-scores of GPL. The trees were reconstructed from observed *E. coli* genome sequences, applying CGP, RAxML, ClonalFrameML, and Gubbins to nucleotide sequences, and CGP and RAxML to amino acid sequences.

## DISCUSSION

We introduced a coarse-graining phylogenetic (CGP) model, which infers a phylogenetic tree from the estimated pairwise coalescent times of genomes. We conducted extensive analyses to compare the accuracy of the CGP algorithm with other state-of-the-art algorithms to demonstrate its ability to reliably predict the topology, branch lengths, and root positioning of phylogenetic trees. The CGP model does not rely on the assumption of low recombination rates, which allows it to predict branch lengths accurately even if the vast majority of the considered genome segments have experienced recombination on the timescale covered by the phylogeny.

Analyses performed on the real *E. coli* genome sequences showed that trees reconstructed from core genome amino acid sequences are more accurate, *i.e.*, more consistent with the signal inferred from the distribution of genes in the extant genomes, than trees calculated from nucleotide sequences. Amino acid sequences of core genes tend to evolve more slowly than the corresponding DNA sequences, as these genes show dN/dS values <1 [28], and accordingly, the divergence of a pair of amino acid sequences is lower than that of their nucleotide counterparts (see Supplementary File S3 for nucleotide and amino acid sequence divergence between *E. coli* genome pairs). Thus, amino acid sequences may be more “clonal” than nucleotide sequences and thus may provide more accurate phylogenetic signals.

The computational demand of the CGP algorithm is independent of sequence length, as CGP considers only SSP distributions that are represented by a vector of 1+*l*_*s*_^*cutoff*^ elements in the computer code. Calculation of the CGP score (Eq. (3)) involves multiplication of (1+*l*_*s*_^*cutoff*^)×(1+*l*_*s*_^*cutoff*^) matrices; thus, the computational time scales as *O*((*l*_*s*_^*cutoff*^)^*k*^), where *k*≤3 depends on the algorithm that carries out the matrix operations. When reconstructing a tree of *n* genomes, the score calculation involves the summation over *n*(*n*-1)/2 pairs, making it scale as *O*(*n*^2^). The segment size *ls* affects the efficiency and accuracy of the algorithm. While a smaller *l*_*s*_ leads to lower accuracy, increasing *l*_*s*_ leads to higher computational demand; a large *l*_*s*_ combined with a small *l*_*s*_^*cutoff*^ can also reduce the accuracy. Hence, one needs to set *l*_*s*_ and *lscutoff* carefully to balance the need for speed and accuracy.

The current algorithm that implements the CGP model is very simple; it should be considered a proof of concept. While it makes use of Monte Carlo simulation to sample the tree+parameter space, a hill-climbing method may be more efficient. Other possible improvements involve better local search moves in the ultrametric tree space; one might even drop the stringent ultrametricity constraint, and replace it with a more flexible matrix-tree mapping method that allows a more efficient search in the tree space. The mutation matrix in the current model can be improved to include back mutations and a more complex mutation model. We leave these possible improvements to future studies.

## ACKNOWLEDGEMENT

We would like to thank Martin Lercher for helpful comments and advice. This work is supported by the German Research Foundation (DFG-grant CRC 680 awarded to Martin Lercher).

## SUPPORTING INFORMATION

1. Source code of the CGP algorithm: https://github.com/TinPang/coarse-graining-phylogenetics
2. Supplementary Text, Figures and Tables.
3. Supplementary File S1: Analyses of the simulated genomes: the symmetric distance (SD) and branch score distance (BSD) between the reconstructed trees and the true trees.
4. Supplementary File S2: Analyses of the real genomes: b-number of the *E. coli* core genes used to make the esuper-gene sequences, strains in each test-group, and also the GLOOME posterior likelihood (GPL) of the reconstructed trees.
5. Supplementary File S3: Sequences and their phylogenetic trees: sequences of the simulated genomes in different test-groups, their true trees and also the phylogenetic trees reconstructed by different algorithms; sequences of the *E. coli* genomes and their trees reconstructed by different algorithms; genes to orthologous gene families map provided by ProteinORTHO.

